# Histone modifications analysis reveals enhancers reprogramming during maternal-to-zygotic transition

**DOI:** 10.64898/2026.05.06.723106

**Authors:** Kaiyue Hu, Chunling Wang, Dong Fang, Jiacheng Lu, Xiangrui Meng, Lingling Chen, Yage Yao, Jia Guo, Sarmir Khan, Wenbo Li, Yaqian Wang, Yang li, Hao Chen, Jiawei Xu

## Abstract

Enhancers are important epigenetic regulatory components that coordinate spatiotemporal gene expression and play a crucial role in mammalian development, gene regulation, and illness. During early mammalian embryogenesis, histone modifications including H3K4me1, a canonical enhancer mark, and H3K27ac, which identifies active enhancers, are still poorly understood. This study profiles the genome-wide H3K4me1 and H3K27ac patterns in mouse oocytes and pre-implantation embryos using low-input CUT&RUN (cleavage under targets and release using nuclease) with input as low as 50 cells. Both markers are enriched in distal regions and exhibit distinct sequence preferences and reprogramming dynamics in pre-implantation embryos. H3K27ac, a mark of active enhancers, is reprogrammed at the 2-cell stage, whereas H3K4me1 is remodeled at the 4-cell stage and overlaps with H3K27ac, coinciding with accessible chromatin regions. Interestingly, H3K4me1 and H3K27ac are co-localized in promoter regions, where they are mutually exclusive to H3K4me3. Enhancers are dynamically remodeled during maternal-to-zygotic transition (MZT). there are three enhancer types: active enhancers (H3K4me1/H3K27ac), primed enhancers (H3K4me1), and poised enhancers (H3K4me1/H3K27me3). Active enhancers increase significantly after zygotic genome activation. Further, super-enhancers are present across the genome and are mainly enriched in promoters. The specific motifs enriched by super-enhancers may be associated with the variations of gene expressions at different stages.

## Introduction

Epigenetic regulation, including global DNA demethylation, chromatin remodeling, spatial genome reorganization, and extensive transcriptional changes, is crucial for embryonic development and cell differentiation after fertilization^[1-4]^. One important potential cause of aberrant embryonic development is incomplete reprogramming of epigenetic modifications. Histone modifications, also known as histone marks, are key epigenetic regulators and are essential for the dynamic reprogramming of chromatin architecture, function, and gene expression^[5, 6]^.

Chromatin Immunoprecipitation sequencing (ChIP-seq) has been a potent technique for tracking genome-wide mapping of histone modifications^[7-9]^. Notably, the distinctive landscapes of histone marks throughout early embryonic development have been revealed by the low-input and limited cell number chromatin profiling tools^[10-12]^. It has been reported that H3K4me3 is rapidly erased from the paternal genome after fertilization but re-established during major zygotic genome activation (ZGA). Conversely, the maternal genome retains a non-canonical form of H3K4me3 (ncH3K4me3)^[13-15]^. Moreover, overexpression of the histone demethylase Kdm5b has been shown to reactivate the transcriptome in mature oocytes, suggesting that ncH3K4me3 may drive genome-wide transcriptional silencing during oocyte maturation^[15]^. In contrast to the ncH3K4me3-dependent mode in mice, our earlier study demonstrated that human germinal vesicle (GV) and metaphase I (MI) oocytes lack substantial ncH3K4me3, indicating a species-specific mechanism of gene expression regulation in human oocytes^[16]^. Interestingly, the width of the H3K4me3 peaks is highly dynamic during mouse preimplantation embryonic development and shows a positive association with gene expression levels^[14]^. It has also been demonstrated that other histone modifications, such as H3K27me3, H2AK119ub1, and H3K9me3, are important for early embryogenesis^[17-19]^. Collectively, these studies underscore that histone modifications are key determinants for active and inactive chromatin domains and are essential for chromatin remodeling during oogenesis and the maternal-to-zygotic transition (MZT)^[20]^.

Enhancer elements throughout the genome are frequently identified and annotated using the histone mark H3K4me1^[21, 22]^. Previous studies on H3K4me1 have mostly focused on enhancer activity in adult tissues^[23-25]^ and embryonic stem cells (ESCs)^[26]^. Little is known about the distribution and regulatory mechanisms of H3K4me1 during the early embryonic development in mammalian.

The dynamic role of enhancer marks in early development is shown by evidence from model organisms. H3K27ac is frequently used to distinguish active from inactive enhancers and to predict the developmental states ^[21, 22, 27-29]^. The parental-to-zygotic transition in zebrafish is supported by genome-wide H3K27ac dynamics^[30]^. Similarly, enhancer mapping (H3K27ac) revealed TCF3/12 as critical regulators of folliculogenesis in mouse oocytes and early embryos^[31]^. In *Xenopus tropicalis*, enhancer marks (Ep300, H3K27ac, and H3K4me1) work with maternal transcription factors (Otx1, Vegt, and Foxh1) to regulate endodermal specification during ZGA^[32]^. While recent research has characterized the rapid dynamics of H3K27ac and its role in triggering ZGA, the lack of integrated H3K4me1 profiling leaves the full developmental trajectory and the initial ‘priming’ of these enhancers during the maternal-to-zygotic transition largely unresolved.

In our previous study, we successfully captured H3K27ac reprogramming in 8-cell embryos and inner cell mass (ICM) cells by optimizing CUT & RUN technology to generate high-quality data from as few as 50 human ESCs^[16]^. In this work, we use this technique to generate the first comprehensive, multi-dimensional genome-wide maps of H3K4me1 and H3K27ac in mouse oocytes and early embryos. By integrating these histone modification marks with transcriptomic and chromatin accessibility data, our analysis goes beyond identifying “active” sites; it systematically delineates three distinct functional categories of enhancers—active (H3K4me1/H3K27ac), primed (H3K4me1), and poised (H3K4me1/H3K27me3)—characterizing their unique spatiotemporal dynamics and associated signaling pathways. Ultimately, this study provides the first comprehensive overview of H3K4me1 and H3K27ac reprogramming during oocyte maturation, embryonic development, and cell differentiation, thereby deepening our understanding of how the combinatorial epigenetic code governs the transition from oocyte maturation to embryonic cell fate.

## Results

### 1. H3K4me1 and H3K27ac are widely distributed in mouse oocytes and pre-implantation embryos

To study the histone modification H3K4me1 and H3K27ac in mouse oocytes and pre-implantation embryos, we firstly carried out the immunofluorescence staining to investigate the alteration of H3K4me1 and H3K27ac at each stage (Figure 1A). For H3K4me1, it showed the most notable fluorescence signal intensity in the GV oocyte, while the signal intensity was relatively much weaker in the zygote and 2-cell embryo, followed by an increased signal intensity in the 4-cell to blastocyst stage. Unlike H3K4me1, H3K27ac undergoes erasure in MII oocytes and re-establishment after fertilization. Overall, the histone modifications H3K4me1 and H3K27ac are widespread in preimplantation embryos.

**Figure 1.**
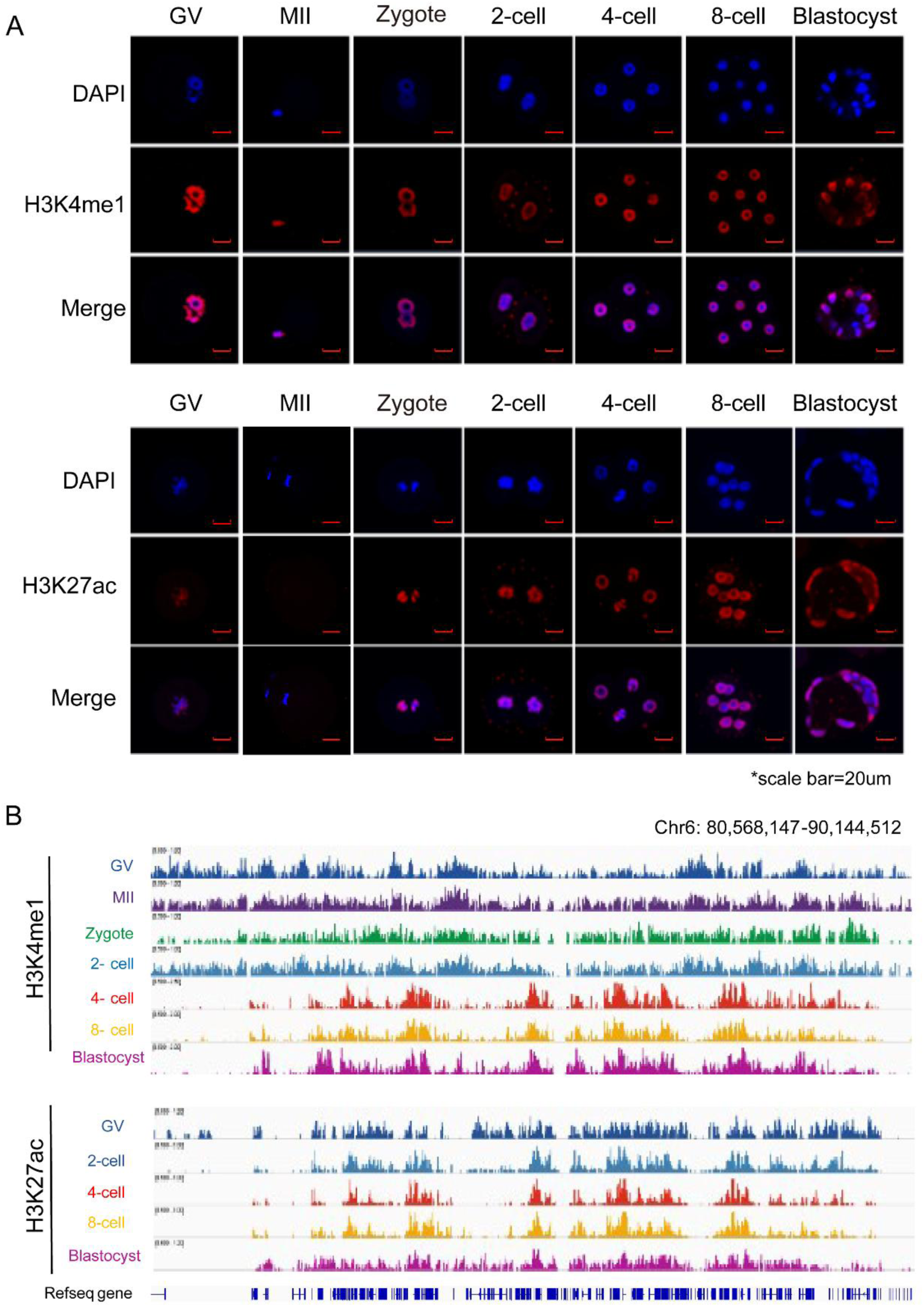
Widespread distribution of H3K4me1 and H3K27ac in mouse oocytes and pre-implantation embryos. A, Immunostaining shows H3K4me1 and H3K27ac signals in GV, MII, Zygote, 2-cell, 4-cell, 8-cell, and blastocyst stages respectively. Scale bar, 20 μm. B, IGV browser view shows H3K4me1 and H3K27ac enrichment in mouse oocytes and pre-implantation embryos.

CUT & RUN is a powerful general-purpose technique for detecting protein-DNA interactions in the cell’s native chromatin environment^[33-35]^. In the previous study, we have validated effectiveness of CUT & RUN to define the genome-wide distribution of H3K27ac in the human embryos^[16]^. Therefore, here, we only validated the effectiveness of CUT & RUN technique in detecting H3K4me1. Our data showed that all the data in ESCs generated from 50 cells to 10 thousand cells was with excellent signal-to-noise ratio (SNR) and the global H3K4me1 enrichment was consistent with the ENCODE data which is from more than 10 thousand cells. Moreover, each group of data was well comparable with data from 10 thousand cells and ENCODE data, with the Pearson’s correlation coefficients between each group of data and 10000 data were all above 0.9, and Pearson’s correlation coefficients between each group of data and Encode data is about 0.8 (Figure S1, A and B). These above suggest the high applicability of CUT & RUN method to capture H3K4me1 and H3K27ac using as low as 50 cells. Then, we profiled H3K4me1 and H3K27ac in mouse oocytes and embryos. Each data has been repeated at least twice, and two replicates for each embryonic stage were deeply well correlated, with the Pearson’s correlation coefficients between each two replicates of H3K4me1 for each above 0.9, and that of H3K27ac were between 0.862 and 0.972, indicating the high duplicability and reliability of the H3K4me1 and H3K27ac data (Figure S1, C and D).

We then analyzed the distribution characteristics of H3K4me1 and H3K27ac in mouse oocytes and pre-implantation embryos. Firstly, our data showed that H3K4me1 and H3K27ac were widely distributed throughout the whole genome (Figure 1B). Secondly, compared with GV oocytes, MII oocytes, zygotes and 2-cell stages, it should be noted that the enriched regions of H3K4me1 after the 4-cell stage were significantly reduced, accompanied by a significant change of the distribution pattern. Nonetheless, the cluster analysis results are not consistent with our expectations as the distribution patterns of each stage before or after 4-cell stage were obviously different, which might be due to the considerably widespread enrichment in the genome (Figure S2A). In particular, the H3K27ac signal erase and the distribution pattern transition occurred between the GV and 2-cell stages, which is earlier than that of H3K4me1. Moreover, the result of cluster analysis revealed that the distribution patterns of H3K27ac were almost identical before or after 2-cell stage (Figure S2B). Together, these data preliminarily confirmed that H3K4me1 and H3K27ac were reprogramming during pre-implantation embryo development.

To further identify the distribution characteristics of H3K4me1 and H3K27ac during early embryonic development, we performed peak calling analysis using model-based analysis (MAC) and analyzed the genomic distribution. The results showed that H3K4me1 and H3K27ac enriched in multiple genomic elements, such as promoter, 5’UTR, 3’UTR, exon, intron, downstream and distal intergenic (Figure S2, C and E). In this study, we also partitioned the genomes into promoter regions (TSS ± 3kb) and distal regions (the site beyond 3kb from TSS), we found both were enriched in the distal regions, which is similar with discoveries reported in the adult tissues^[36, 37]^. In detail, about 90% of H3K4me1 peaks located in the distal regions. And the distribution of H3K4me1 across various genomic elements stabilized from the 4-cell stage onward. Prior to this stage, the percentage ranged from 5.51% to 9.1% in promoter regions and from 90.9% to 94.49% in distal regions. From the 4-cell stage onward, these percentages stabilized at 11.1%–11.64% in promoter regions and 88.36%–88.9% in distal regions (Figure S2D), which indicated the reprogramming of H3K4me1 potentially occurs during oocyte maturation and ZGA stages. Meanwhile, H3K27ac peaks are also predominantly enriched in distal regions, which is consistent with H3K4me1. However, the distribution of H3K27ac altered from 2-cell stage (major ZGA) onwards, with the percentage fluctuating around 10% in the promoter regions before 2-cell stage, whereas the percentage ranging from 15.57% to 18.17% from 2-cell stage to 8-cell stage in the promoter region, suggesting the reprogramming of H3K27ac was dynamic during oocyte and embryo development.

### 2. H3K4me1 and H3K27ac exhibit distinct distribution pattern compared to H3K4me3 at promoter regions during preimplantation development

As mentioned above, we successfully detected the dynamic signals of H3K27ac and H3K4me1 in the promoter regions, therefore, we focused on analyzing the reprogramming and their potential functions in the promoter regions of each oocyte and preimplantation embryo stage. We firstly analyzed the distribution pattern of H3K4me1 and H3K27ac in the promoter regions (TSS ± 3kb) and our results showed that H3K4me1 peaks are enriched in promoters around TSS with a bimodal pattern. Besides, compared with MII oocyte, zygote and 2-cell stage, H3K4me1 signals are more significantly enriched in the promoter regions in GV oocyte, 4-cell, 8-cell and blastocyst stage. For H3K27ac, the signals are mainly enriched in the promoter region in a canonical pattern and the signals in 2-cell, 4-cell and 8-cell stage were more pronounced than those in GV oocyte and blastocyst stage (Figure 2A and Figure S3A).

**Figure 2.**
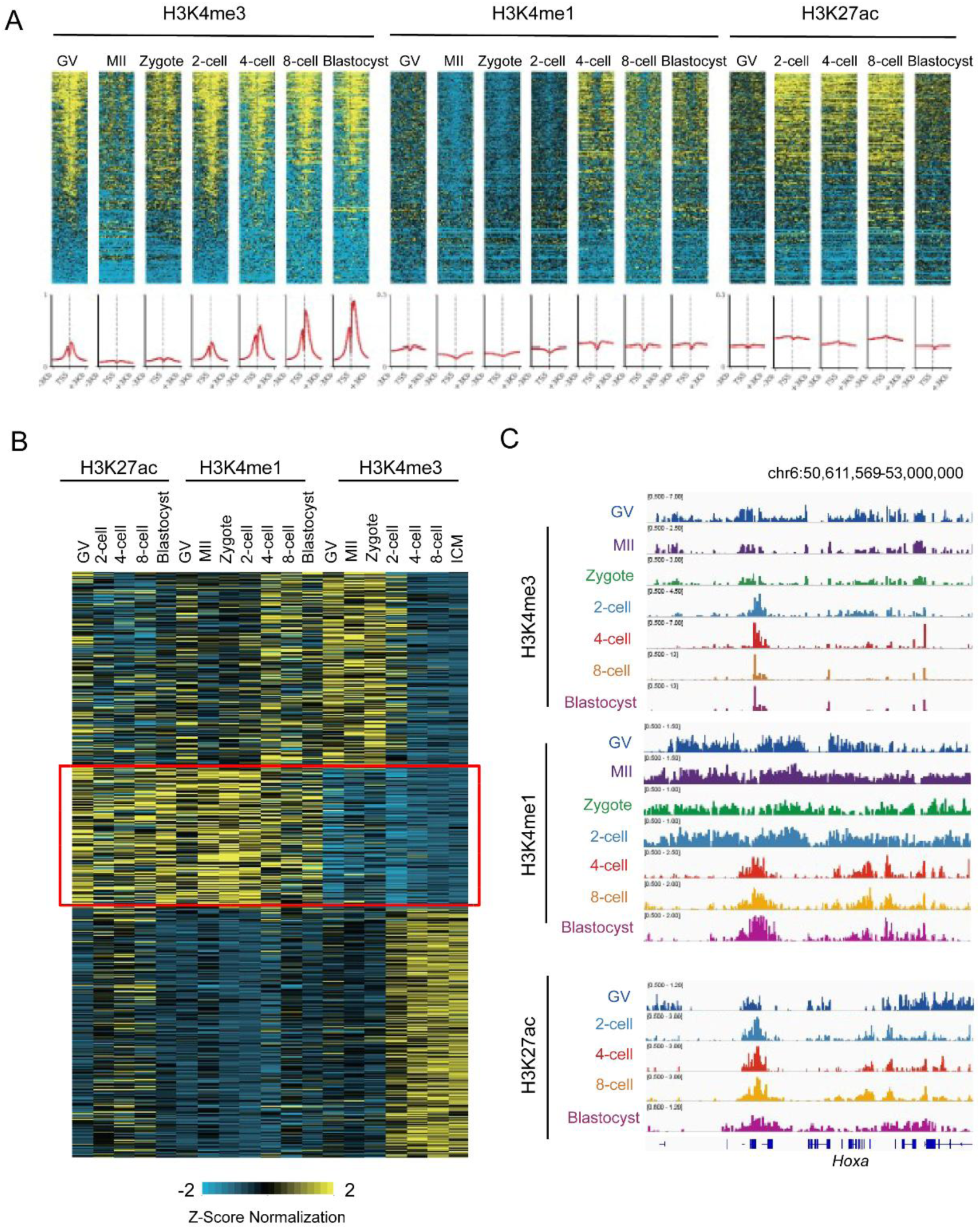
Distinct distribution patterns of H3K4me1 and H3K27ac relative to H3K4me3 at promoter regions. A, Heatmap shows H3K4me3, H3K4me1, and H3K27ac signals in promoter regions. B, K-means clustering analysis of H3K4me3, H3K4me1, and H3K27ac enrichment in promoter regions. C, IGV browser view shows the H3K4me3, H3K4me1, and H3K27ac around Hoxa regions.

Both H3K4me1 and H3K4me3 are methylated modifications of H3K4. Generally, H3K4me3 marks promoters of active genes. In mature mouse oocytes, H3K4me3 is widely distributed at low levels in a non-canonical pattern, which is also found at the zygote and 2-cell stage^[15]^. However, at the late 2-cell stage, non-canonical H3K4me3 is extensively erased, until the 4-cell embryonic stage basically disappeared, then gradually exhibited in the canonical pattern^[13]^. To better understand the H3K4me1 and H3K27ac distribution in the promoter regions, we comprehensively analyzed the distribution of H3K4me3, H3K4me1 and H3K27ac in the promoter regions of each stage. The results showed that the H3K4me3 was most strongly enriched at TSS, however, H3K4me1 signal was weakest at the TSS, which was contrary to H3K4me3, especially after the 4-cell stage. But this trend was not observed in the distribution of H3K27ac (Figure 2A). Second, we performed the k-means cluster analysis on H3K4me1, H3K27ac and H3K4me3 signal in the promoter regions. It was clearly observed that in the regions marked a strong H3K4me3 signal, the H3K4me1 and H3K27ac signals are relatively weaker. Likewise, both H3K4me1 and H3K27ac can also be observed in regions where H3K4me3 signal is weak, mainly at the pre-ZGA stage (Figure 2B). Third, we analyzed the relationship between the three histone modifications and CpG density. The results showed that the H3K4me3 signal and CpG density signal were significantly weaker in the regions where with stronger H3K4me1 signal. In contrast, H3K4me3 signal and CpG density signal were significantly stronger in regions with weaker H3K4me1 signal. H3K27ac showed the same trend as H3K4me1 (Figure S3B). Finally, the tracks of three kind histone modification in the IGV browser were also presented. Surprisingly, a high degree of overlap was noticed between the H3K4me3 and H3K27ac in the promoters’ regions while the H3K4me1 located on the left and right sides of H3K4me3 and H3K27ac, mainly at the pre-ZGA stage (Figure 2C).

Although the distribution pattern of H3K4me1 in the promoter region tends to be relative stable after ZGA, the pattern of each stage about H3K4me3 and H3K27ac is not the same. Here, we identified the stage-specific peaks and subsequently performed a signal pathway analysis using the gene list enriched in the peaks. Notably, for H3K4me1, the specific genes in GV were involved chromatin organization, in MII were associated in the detection of chemical stimulus involved in sensory perception of bitter taste, genes in zygote related to cellular response to energy homeostasis and genes in blastocyst related to protein modification (Figure S3, C and D). For H3K27ac, stage-specific peaks were enriched in distinct biological processes across developmental stages. Specifically, stage-specific genes in the 4-cell stage were highly enriched in the cellular response to DNA damage stimulus, while those in the GV, 2-cell, 8-cell, and blastocyst stages were significantly associated with processes related to the regulation of fatty acid biosynthetic process, DNA alkylation, positive regulation of programmed cell death, and regulation of the Wnt signaling pathway, respectively (Figure S3, F and G). In addition, the distribution of H3K4me1 or H3K27ac at several stage-specific loci were shown out, respectively (Figure S3, E and H).

### 3. Reprogramming of H3K4me1 and H3K27ac in distal regions during preimplantation development

Our results have demonstrated that H3K4me1 and H3K27ac were extremely widespread (almost 90%) in distal regions of mouse oocytes and preimplantation embryos. Therefore, we specifically analyzed the distribution of H3K4me1 and H3K27ac in distal regions and explored their reprogramming to deduce their possible mechanisms involved in oocyte maturation and embryonic development. First, we identified the number of H3K4me1 and H3K27ac peaks in the distal regions of each stage. For H3K4me1 peaks, the numbers identified at each stage were 592809 in GV oocyte, 874601 in MII oocyte, 847315 in zygote, 633109 in 2-cell, 486976 in 4-cell, 446104 in 8-cell and 481616 in blastocyst (Figure 3A). As shown in Figure 3B, the number of H3K27ac peaks identified at each stage was 33780 in GV oocytes, 37777 in 2 cells, 37958 in 4 cells, and 49332 in 8 cells and 24184 in blastocysts. Based on the number of identified peaks at each stage, it is concluded that the peaks of H3K4me1 and H3K27ac were not stably inherited from the oocyte to the preimplantation embryo, but constantly underwent changes. H3K4me1 peaks decreased significantly from the MII stage to the 4-cell stage but fluctuated in a small range from 4-cell to 8-cell stage, suggesting that the H3K4me1 markers in the distal region were continuously reprogramming after fertilization and then became stable at 4-cell stage. Meanwhile, H3K27ac markers increased significantly from GV to 8-cell stage, and then decreased from 8-cell to blastocyst stage (Figure 3B), indicating that the H3K27ac markers in the distal region were erased after 8-cell stage and underwent remodeling from GV to blastocyst stage.

**Figure 3.**
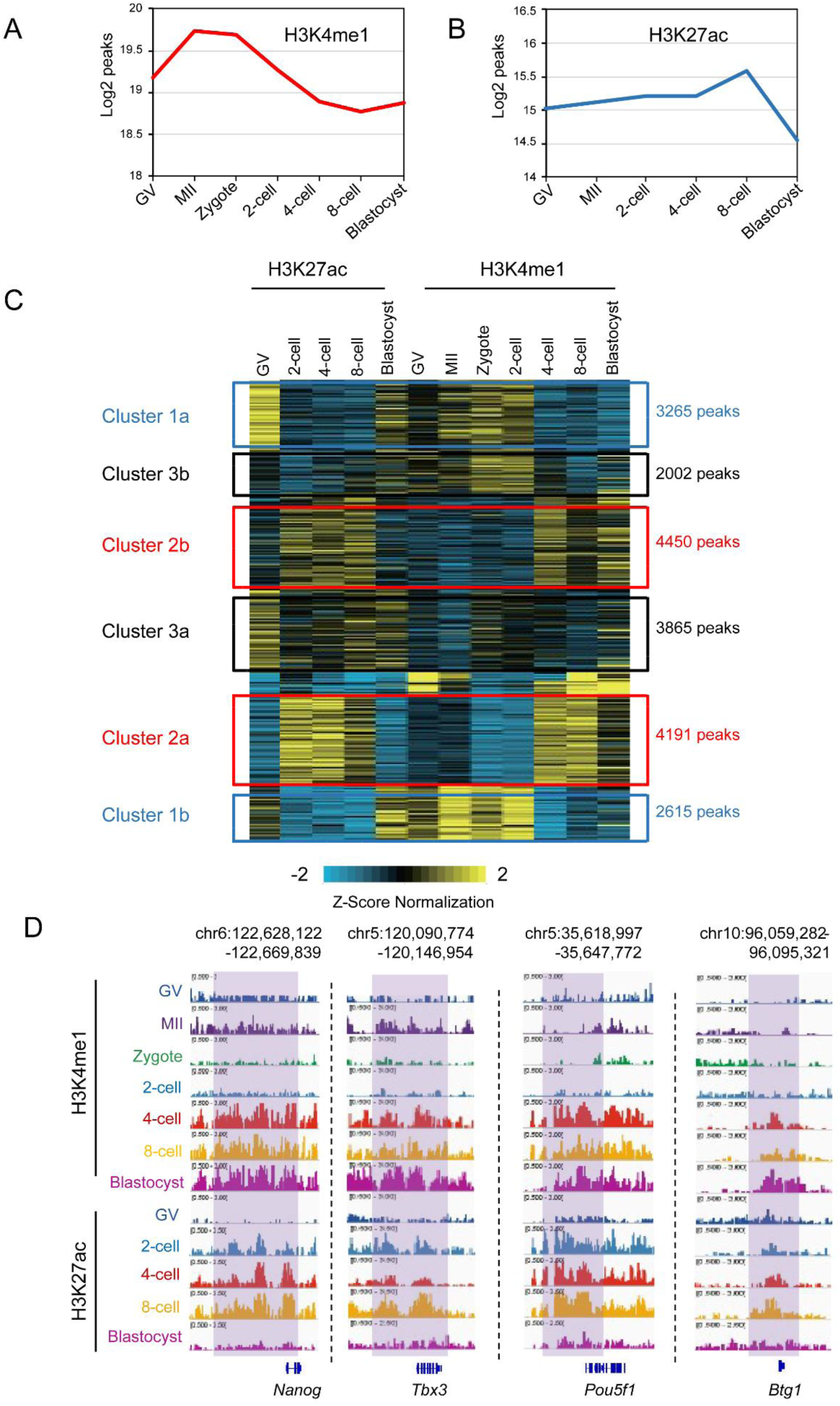
The reprogramming of H3K4me1 and H3K27ac in distal regions. A, The H3K4me1 peaks identified in distal regions. B, The H3K27ac peaks identified in distal regions. C, The reprogramming of co-location H3K4me1and H3K27ac in GV oocytes within distal regions. D, IGV browser view showing H3K4me1 and H3K27ac enrichment at sites of totipotent factors.

In order to understand the co-localization of H3K4me1 and H3K27ac peaks in distal regions, K-means clustering analysis was performed. The resulting heatmap revealed distinct reprogramming dynamics: H3K27ac signals were established as early as the 2-cell stage and maintained through the 8-cell stage, whereas H3K4me1 peaks underwent extensive resetting and were established from the 4-cell stage. Accordingly, as shown in Cluster 1, pre-existing H3K27ac marks extensively co-localized with newly established H3K4me1 peaks, specifically from the 4-cell and 8-cell stages (Figure S4A). Meanwhile, the hierarchical cluster analysis of H3K4me1 and H3K27ac also support these results (Figure S4B). Next, to gain a deeper understanding of H3K4me1 and H3K27ac reprogramming, we focused on those peaks of H3K4me1 and H3K27ac which are inherited, erased. Considering the importance of maternal materials for embryonic development and the fact that the number of H3K4me1 and H3K27ac peaks were reprogrammed from GV oocytes. Hence, we firstly focused on the reprogramming of H3K27ac and H3K4me1 in GV oocytes. It was obvious that the H3K27ac peaks in GV oocytes were mainly enriched in cluster 1a and cluster 3a, both were erased in 2-cell stage and 4-cell stage, the cluster 1a was re-built in blastocyst while the H3K4me1 cluster 1a was significant different from H3K27ac, the cluster 3a was rebuilt in 8-cell and blastocyst stage, while the H3K4me1 cluster 3a seemed low across every stage. In addition, domains with no apparent H3K27ac signal in GV oocytes were classified as cluster 2, and interestingly, no H3K4me1 signal was detected in these domains either. However, the H3K27ac and H3K4me1 were further established in the corresponding regions at 2-cell and 4-cell respectively, co-localized in 4-cell stage, and stably inherited to blastocyst. Meanwhile, these results confirmed that in the distal regions, the reprogramming of H3K4me1 occurred at 4-cell stage and the reprogramming of H3K27ac occurred at 2-cell stage (Figure 3C). Notably, from 4-cell to blastocyst stage, strong H3K4me1 and H3K27ac signals were detected near some totipotent factors, such as *Nanog, Tbx3, Pou5f1* and *Btg* (Figure 3D).

Extensive erasure of H3K4me3 in the distal region after ZGA was demonstrated to be associated with the activation of the H3K4me3-related demethylase Kdm5b^[38]^. Considering that histone H3 lysine 4 (H3K4) could be mono-, di-, and tri-methylated, we therefore analyzed the expression levels of mRNAs for all H3K4-related demethylases and methyltransferases in oocytes and early embryos to explore whether reprogramming of H3K4me1 is driven by a key enzyme. The transcriptomic data revealed that among these enzymes, the demethylases Kdm1b and Kdm5b exhibited the most prominent, yet temporally distinct, expression patterns (Figure S4, C and D). Specifically, Kdm5b expression surged from ZGA, whereas Kdm1b was highly expressed in GV oocytes and progressively decreased until the 4-cell stage. Since Kdm5b acts primarily on H3K4me3, we asked whether the dynamic changes in Kdm1b expression could affect H3K4me1 generation. Notably, the continuous decline in Kdm1b expression perfectly coincided with the extensive erasure of H3K4me1 peaks. Furthermore, previous reports proposed that Kdm1b acts primarily on H3K4me2^[39-41]^, suggesting that the extensive erasure of H3K4me1 peak from the oocyte to the 4-cell stage may be associated with a progressive decrease in the expression level of kdm1b, however, which core factors are directly responsible for the reprogramming of H3K4me1 still need to be further explored.

### 4. Distal H3K4me1 and H3K27ac peaks highly overlapped with chromatin accessibility region

Chromatin plasticity is critical for regulating essential cellular processes such as DNA replication, repair, and transcription^[42, 43]^. Specific histone modifications are instrumental in the spatiotemporal regulation of gene expression and the modulation of promoter activity. Therefore, we aimed to explore the relationship between the chromatin accessibility landscape and histone marks (H3K4me1 and H3K27ac) in preimplantation embryos.

We firstly global view the genome-wide distribution of H3K4me1, H3K27ac, and chromatin accessibility. To our interesting, we observed that both the H3K4me1 and H3K27ac peaks were highly overlapped with the ATAC-seq signals from 4-cell stage onwards (Figure 4A and Figure S5A), indicated the critical regulatory role of H3K4me1 and H3K27ac on gene expression.

**Figure 4.**
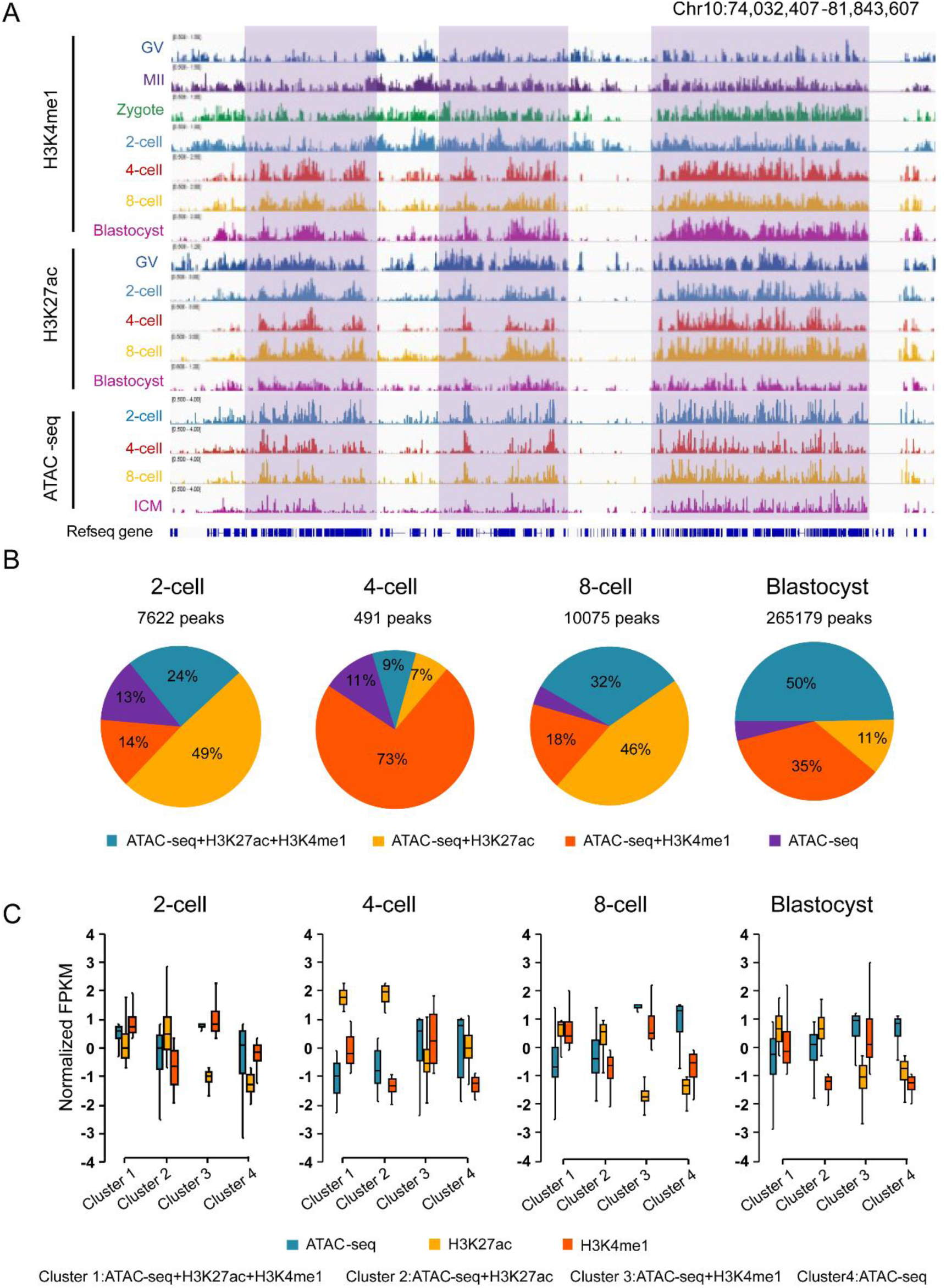
H3K4me1 and H3K27ac highly overlap with open chromatin regions. A, IGV browser view showing H3K4me1 and H3K27ac signals highly overlapping with open chromatin regions. B, Pie chart showing the overlap among H3K4me1, H3K27ac and ATAC-seq peaks. C, Boxplot displaying normalized FPKM values for H3K4me1, H3K27ac and ATAC-seq across each cluster.

In order to gain insight in the relationship of H3K4me1, H3K27ac and chromatin accessibility in more detail, next we compared genome-wide peaks distribution of H3K4me1, H3K27ac and ATAC-seq signals. As is shown in Figure 4B, 7622, 491, 10075 and 265179 peaks were respectively identified in the 2-cell, 4-cell, 8-cell and blastocyst stage. Then ATAC-seq peaks in each stage were classified into four clusters based on whether the peaks of H3K4me1 and/or H3K27ac overlap with those of ATAC-seq. That ATAC-seq peaks overlap both H3K27ac and H3K4me1 peaks was considered as cluster 1, ATAC-seq peaks overlap H3K27ac peaks only was considered as cluster 2, ATAC-seq peaks and H3K4me1 peaks only was considered as cluster 3 and ATAC-seq peaks without H3K27ac and/or H3K4me1 peaks was considered as cluster 4. Meanwhile, normalized FPKM of H3K4me1, H3K27ac and ATAC-seq signals of each cluster were calculated and displayed in Figure 4C. It is clear that the vast majority of ATAC-seq peaks (87%∼92%) in each stage were overlapped by H3K4me1 and/or H3K27ac peaks to varying degrees, further implying the critical regulatory role of H3K4me1 and H3K27ac in the region of chromatin accessibility in post-ZGA. Additionally, compared to that in 2 cell stage, ATAC-seq peaks were predominantly overlapped by H3K4me1 signals in the 4-cell stage, which is consistent with our previously mentioned result that the reprogramming of H3K4me1 in the 4-cell stage. We also observed a gradual increase in the proportion of H3K4me1 and H3K27ac peaks overlapping with ATAC-seq peaks (cluster 1) from 4cell stage to blastocyst, suggesting that colocalization of H3K4me1 and H3K27ac continuously promotes the development of the totipotent embryos.

Furthermore, we analyzed the correlation between H3K4me1 and chromatin accessibility based on both the promoter and distal regions, respectively. Surprisingly, for both H3K4me1 and H3K27ac, our data showed that the correlation coefficient between distal regions and chromatin accessibility were significantly higher than those between promoter regions and chromatin accessibility.

As shown in Figure S5A, the correlation coefficient between H3K4me1 and ATAC-seq in the promoter region of each stage was about 0.3, while in the distal region, the correlation coefficient between the two signals from 2 cells to blastocyst stage was 0.027, 0.928, 0.985 and 0.898. It can be clearly seen that the correlation coefficient between H3K4me1 and chromatin accessibility increases significantly after 4 cells, which is related to the reprogramming of H3K4me1 at the 4-cell stage. Similar to H3K4me1, the correlation coefficient between H3K27ac and ATAC-seq signals in the promoter region was around 0.4 (0.425 in 2-cell, 0.425 in 4-cell, 0.425 in 8-cell, 0.420 in blastocyst), while in the distal region, the correlation coefficients between H3K27ac and ATAC-seq signals were all higher than 0.9 (0.938 in 2-cell, 0.937 in 4-cell, 0.959 in 8-cell, 0.951 in blastocyst) (Figure S5B). Therefore, correlation analysis of H3K4me1 and H3K27ac with ATAC-seq data showed that H3K4me1 and H3K27ac signaling and chromatin accessibility are highly overlapping in distal regions.

### 5. The transition of typical enhancer during preimplantation development

Enhancers are distal regulatory elements that can activate time and spatial specific gene expression^[44]^. In mammalian, enhancers are usually marked by H3K4me1, and H3K27ac and H3K27me3 can further functionally classify enhancers marked by H3K4me1^[22]^. Then, we further investigated the distribution and characteristics of different types of enhancers and their roles during preimplantation embryonic development. Consistent with previous studies^[45]^, we integrated H3K4me1, H3K27ac and published H3K27me3 data (GSM2041069)^[46]^ to classify all the enhancers into three groups, active enhancers (H3K4me1, H3K27ac), primed enhancers (H3K4me1), and poised enhancers (H3K4me1, H3K27me3) and presented the distribution of the three classes of enhancers in distal regions at different stages (Figure 5, A and B). It could be clearly found that the distribution of the three enhancers was constantly and rapidly changing during preimplantation embryonic development. In oocyte, active enhancers were significantly more than that of inactive enhancers, however, in the 2-cell stage, we found a marked decrease of the active enhancers, followed by an increase in the 4-cell and 8-cell stage, and finally decreased again in blastocyst stage (Figure 5C). Second, we conducted the GO signal pathway analysis to explore the potential functions of three classes of enhancer during the embryo development. The results indicated that active enhancers were mainly enriched in the pathways such as GTPase binding, GTPase activator activity, nucleoside-triphosphatase regulator activity, and DNA-binding transcription factor binding. Primed enhancers were mainly enriched in the pathways such as channel activity, passive transmembrane transporter activity, and metal ion transmembrane transporter activity. Poised enhancers were mainly enriched in the pathways such as cell adhesion molecule binding, pheromone receptor activity, gated channel activity, and DNA-binding transcription activator activity (table S2-S7). Finally, we confirmed the distribution characteristics of histone modifications in three classes of enhancers (Figure 5D). It could be noticed that H3K27ac levels were significantly higher in active enhancers (H3K4me1, H3K27ac) than that in the other two classes of enhancers, primed enhancer (H3K4me1) and poised enhancers (H3K4me1, H3K27me3). Also, the H3K27me3 levels were significantly higher in poised enhancers than that in the active enhancer and primed enhancers. Although H3K4me1 could be enriched in the three classes of enhancers, the H3K4me1 level was highest in active enhancers and lowest in poised enhancers in each stage.

**Figure 5.**
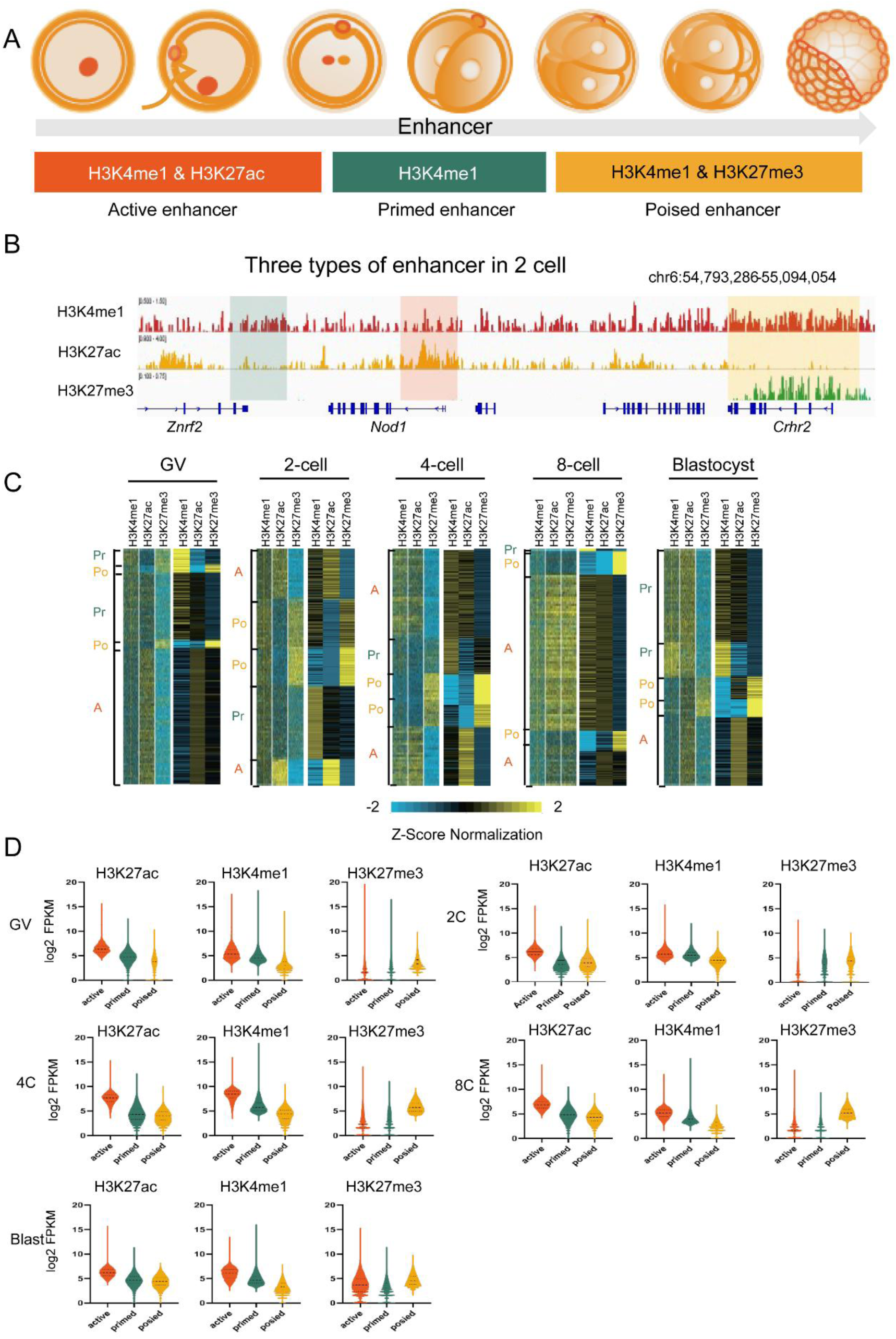
Classification and dynamic landscape of representative enhancers. A, Model of three types of enhancers. B, Three types of enhancers in 2 cell stage. The primed enhancer, active enhancer and posied enhancer were respectively showed in the geen, orange and yellow region. C, Heatmap of H3K4me1-bound enhancers generated by k-means cluster analysis. Each window represents signal 5 kb of the H3K4me1 peak midpoint. Active enhancers are designated A, the primed enhancers are designated Pr, and the poised enhancers are designated Po. D, Violin diagram showing the H3K4me1, H3K27ac, and H3K27me3 distribution in three types of enhancers.

### 6. Identification and function analysis of super enhancers during preimplantation development

Super enhancers differ from typical enhancers in size, distribution, density and content of transcription factor, ability to activate transcription, and sensitivity to interference^[47, 48]^. These enhancer domains are occupied by master transcription factors and associate with genes encoding key regulators of cellular identity, with greater functional specificity^[48]^. The identification of super-enhancers was analyzed using H3K27ac data and it could be seen that the expression levels of the genes *Cnbp* and *Hic2* regulated by super-enhancers were significantly increased at 4-8 cell stage (Figure 6A). In addition, the genomic distributions of super-enhancers were quite different from typical enhancers in the post-ZGA stage, we observed that the region occupied by the super-enhancer is mainly the promoter regions, but not the distal intergenic regions, indicating the potentially efficient interaction between super-enhancers and promoters due to close proximity or overlap^[49]^ (Figure 6B).

**Figure 6.**
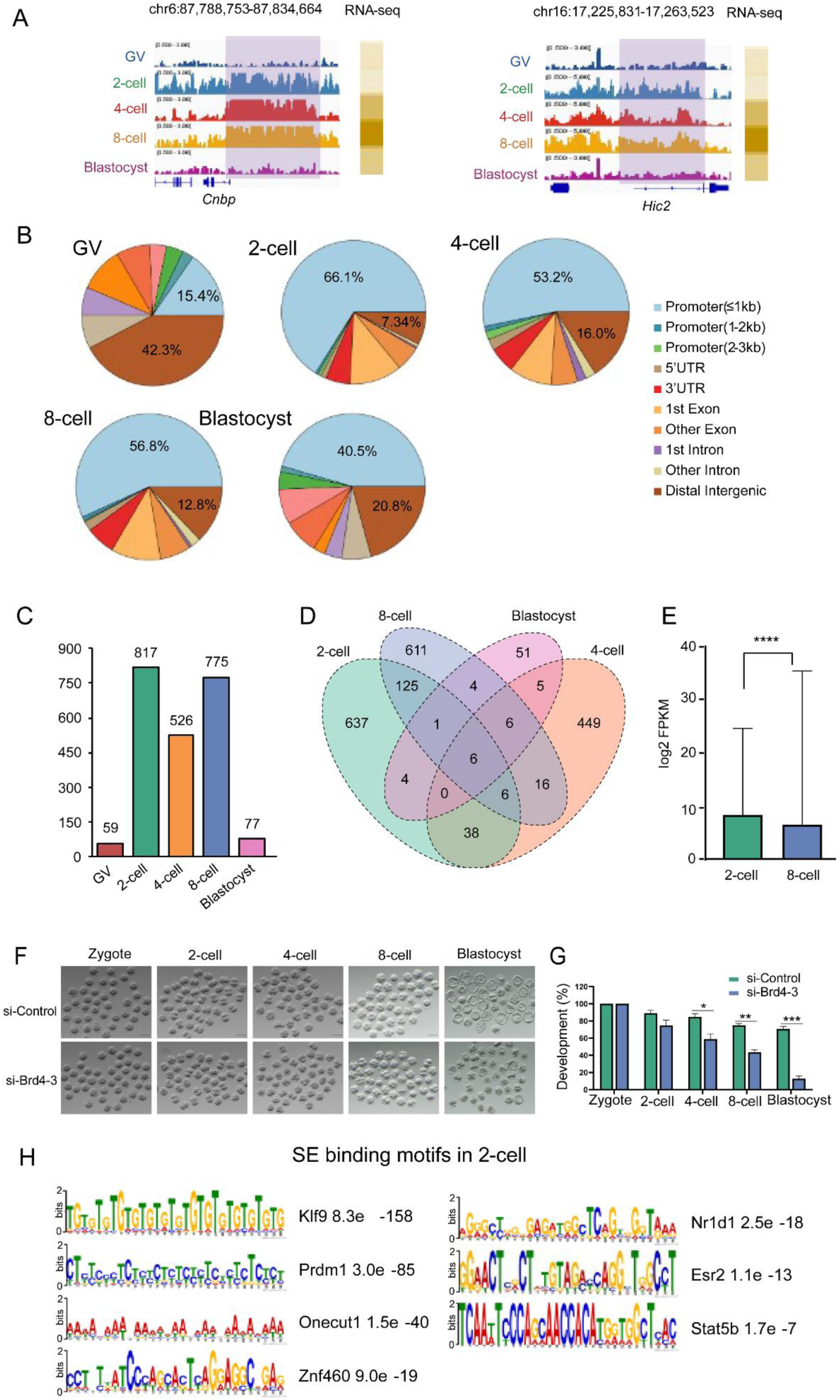
Identification and functional validation of super enhancers. A, The representative loci with the super-enhancer distributed. B, The distribution of the super-enhancer identified in each stage. C, The number of super-enhancers identified in each stage. D, Veen diagram showing the overlap of the super-enhancer identified in each stage. E, The average expression level of genes regulated by the super-enhancers identified in 2-cell and 8-cell stage. F, Representative images of embryos at different developmental stages with si-Control or si-Brd4-3. G, The developmental rates of embryos in si-Control and si-Brd4-3 groups. H, Super-enhancer binding motif identified in 2-cell stage. Data are presented as mean ± SEM. \**P*< 0.05, \*\**P*< 0.01, \*\*\**P*< 0.001.

Next, we further explored the possible regulatory role of super-enhancers on gene expression. Firstly, we system compared the super-enhancers across different stages, interestingly, we found that the number of super-enhancers increased significantly after the 2-cell stage, followed by a sharp decrease in the blastocyst stage. Moreover, 2-cell and 8-cell stages harbor and shared the most super-enhancers (Figure 6, C and D). Therefore, we compared the expression level of genes regulated by super-enhancers shared at the 2-cell and 8-cell stages to explore the potential difference effect on the gene expression regulated by the same enhancer in different stages. Our results indicated that the average gene expression level of 2-cell was significantly higher than that of 8-cells, suggesting that 2-cell-specific super-enhancers had a stronger effect on gene expression activation (Figure 6E). To further functionally validate our sequencing results and confirm the importance of super-enhancers at the 2-cell stage, we targeted BRD4, a key transcription factor and critical component of super-enhancers. We first screened three different siRNAs targeting Brd4 in embryonic stem cells and identified si-Brd4-3 as the most effective one, which significantly reduced Brd4 mRNA levels (Figure S6A). Subsequently, by knocking down Brd4 with si-Brd4-3 to impair super-enhancer function in embryos, we observed that embryo development was severely arrested at the 2-cell stage. Consequently, the developmental rates of embryos reaching the 4-cell, 8-cell, and blastocyst stages were significantly decreased compared to the si-Control group (Figure 6, F and G). This biological phenotype strongly corroborates our sequencing results, demonstrating that super-enhancers play a crucial role during the 2-cell to blastocyst stage. Furthermore, the MEME motif analysis was used to identify the binding motifs for super enhancer in 2-cell. The top seven highly significant predicted motifs were showed in Figure 6H, including *Klf9, Prdm1, Onecut1, Znf460, Nr1d1, Esr2,* and *Stat5b.* Notably, KLF factors are well-known to have critical roles in naive pluripotency^[50]^. *Prdm1* is related to promoter-specific chromatin binding activity^[51]^. Moreover, *Onecut1* is a key factor that enables DNA-binding transcription factor activity and RNA polymerase II cis-regulatory region sequence-specific DNA binding activity^[52]^. Interestingly, all these factors’ motifs were not detected in 8-cell stage, indicating the same super enhancers may play distinct roles throughout development.

## Discussion

Preimplantation embryo development is a highly dynamic and complex process governed by tightly orchestrated genetic and epigenetic regulatory mechanisms^[53, 54]^. The acquisition of totipotency and subsequent embryonic development depend on the considerable epigenetic reprogramming that occurs in the parental genomes after fertilization^[55, 56]^. To avoid developmental abnormalities or embryonic arrest, accurate reprogramming is essential^[57]^. Important histone changes including H3K4me3, which is linked to transcriptional activation, and H3K27me3, which is linked to repression, have been linked to early embryogenesis in both humans and mice, according to earlier research^[58]^. Enhancers are crucial in determining the spatiotemporal patterns of gene expression during development and illness because of their great sequence diversity, tissue specificity, and mechanistic adaptability^[59]^.

In this study, we systematically profiled the genome-wide distributions of the enhancer-associated marks H3K4me1 and H3K27ac throughout mouse preimplantation development, compare the distribution multiple histone modifications and chromatin accessibility, and identify distinct functional types of enhancers by using low-input CUT&RUN technology^[34, 35]^, providing new insights into understanding the regulatory mechanism of early embryonic development and cell fate determination.

After successfully establishing the high-resolution genome-wide map of H3K4me1 and H3K27ac in mouse oocytes and preimplantation embryos, we examined the reprogramming of H3K4me1 and H3K27ac in promoter and distal genomic regions separately. We discovered that reprogramming of H3K4me1 occurs at the 4-cell stage, while reprogramming of H3K27ac occurs at the 2-cell stage. Integrating with the published data of H3K4me3, we found a very interesting phenomenon that H3K4me1 and H3K4me3 show a hide-and-seek relationship throughout the genome. Meanwhile, we also found that the signal distribution of H3K4me1 and H3K27ac in distal regions exhibited substantial overlap with open chromatin regions. On the other hand, to functionally characterize enhancers, we integrated the published H3K27me3 data and classified enhancers into three types—active, primed and poised enhancers—at each developmental stage. Pathway enrichment analysis based on KEGG was conducted to infer their biological roles. We also identified super-enhancers in different stages, and focused on motif difference enriched by enhancers shared in 2-cell and 8-cell stages.

Epigenetic regulation of chromatin, including DNA methylation and histone modifications, are known to be major regulators of ZGA^[60-63]^. Most of the usual epigenetic markers are reprogrammed at 2-cell stage to participated in the ZGA. So, it was worth noting that H3K4me1 is a unique histone modification, our investigation revealed its reprogramming did not occur at 2-cell stage, but at the post-ZGA stage, implying its specific functions mainly involved in the later embryonic development and differentiation. Meanwhile, the revelation of the reprogramming pattern of H3K4me1 will also provide a new perspective for our understanding of the landscape patterns and functions of epigenetic modifications during preimplantation embryonic development H3K4me3 is a well-established gene activation maker, which usually located in promoter regions. Interestingly, previous studies proposed that if H3K4me1 was distributed on both sides of H3K4me3, it could predict a promoter with an active transcriptional state, which was validated in mouse skeletal muscle cells, liver, and pancreatic cells^[64, 65]^. In this study, we found that H3K4me1 co-localized with H3K27ac and exhibited a playing a hide-and-seek with H3K4me3 in the promoter region (Figure 7), especially from the 4-cell to blastocyst stage. This is consistent with the fact that an active transcriptional environment is necessary for the embryo to continue to divide and differentiate after ZGA. Thus, the pattern of H3K4me3 co-localization with H3K4me1 to mark active promoters is not only found in well-differentiated adult tissue cells, but may also be present in pluripotent embryos. However, whether the absence of H3K4me1 affects the function of H3K4me3 needs further study.

In the distal regions, we demonstrated that H3K27ac was reset at 2-cell stage, while H3K4me1 was remodeled at 4-cell stage (Figure 7). Why the timing of the reprogramming is different between H3K4me1 and H3K27ac? With respect to the reprogramming of H3K27ac, we proposed a hypothetical model that H3K27ac needs preferentially to be loaded into active enhancer regions to participate in regulating ZGA, which was further confirmed by super-enhancers identified in 2-cell stage. However, a large number of genomic elements were involved in zygotic genome activation and numerous transcriptional events occurred. Therefore, the absence of H3K4me1 co-localization with H3K27ac at 2-cell stage might avoid the zygotic genome to be hyper-activated to induce the genomic instability. On the other hand, H3K4me1 was extensively widespread in the distal regions, with the number of H3K4me1 peaks in each stage about 10 times more than that of H3K27ac peaks. Besides, the H3K4me1 could function as poised, primed and active enhancer when combined with different modifications, therefore, H3K4me1 enables to balance gene expression across the genome. Indeed, it could be found that active enhancers have the lowest proportion of the three enhancers in the 2-cell stage (Figure 5B), but increased in the 4-cell stage, which also suggested that H3K4me1 was dynamically involved in regulating the global development of the embryo.

Finally, we observed strong overlap between H3K4me1/H3K27ac and open chromatin regions. Thus, we speculated that in the regions enriched in the H3K4me1 and H3K27ac, a large number of cis-regulatory sequences and transposable elements might be located, which is similar to that in the accessible chromatin regions^[66]^. In addition, the combination of H3K4me1, H3K27ac, and accessible chromatin regions could serve as a method to accurately predict transcription factors that play a key role in embryonic development. Taken together, our findings revealed the dynamic engagement of enhancers H3K4me1 and H3K27ac in mouse implantation embryos, and explained the regulatory roles of enhancers and super-enhancers on gene expression. Meanwhile, we proposed a potential role for H3K4me1 in 4 cell-specific reprogramming patterns during embryonic development. Furthermore, we provided a potential approach for predicting key transcription factors. Although there has been a previous study on H3K27ac in mouse embryonic development, this study has some limitations^[13]^. First, it only focused on oocytes, 2-cell and 8-cell stages, and did not include the H3K27ac data of other stage in the preimplantation embryos, so the content of H3K27ac in the study is lacking systematic. Secondly, the study did not fully explain the regulatory mechanism of enhancers on gene expression, and mainly expounds the positional relationship between enhancers and enhancers, as well as enhancers and ZGA genes in different stages. Therefore, how the enhancer marker H3K4me1 and H3K27ac regulate mammalian embryonic development is unclear.

All together, these data lay the foundation for future studies of epigenetic regulatory mechanisms during embryonic development.

## Materials and methods

### 1. Experimental reagents

The details could be obtained in the table S1.

### 2. Oocyte and embryo collection

SPF mice used in the experiment were purchased from Charles River Company (Beijing, China) and raised in the Laboratory Animal Research Center of Zhengzhou University and Henan Academy of Innovations in Medical Science. All animal experiments were approved by the experimental animal center and carried out in accordance with the guidelines for the care and use of laboratory animals. In this experiment, C57BL/6N female mice aged 4-5 weeks and DBA /2 male mice with sexual maturity over 8 weeks were used. 48 hours after PMSG (10 IU) (Solarbio, P9970) injection, female mice were then injected with HCG (10 IU) (Merck Serono S.p.A) and mated with male mice. GV oocytes were obtained 48h after PMSG injection without HCG injection, MII oocytes were obtained at 12-14 hours after HCG injection without mating, and embryos at each stage were obtained according to HCG injection time, such as zygote at 28-30h, late 2-cell at 45-48h, 4-cell at 54-58h, 8-cell at 60-64h, and blastocyst at 84-96h. All the oocytes and embryo were collected and kept in M2 medium (Sigma, M7167). The zona pellucida was medium removed using glutathione solution (Solarbio, G8180) and oocytes and embryo were washed in M2 medium.

### 3. Ethics statement

This study was approved by the medical ethical committee of the First Affiliated Hospital of Zhengzhou University (Approval number 2022-KY-1347) in accordance with the Laws and Regulations of the People’s Republic of China.

### 4. Cell culture

R1 mouse embryonic stem cells (mESCs) were cultured in gelatin-precoated (0.1%, Sigma-Aldrich G1890, ≥5 min) 6-well plates using mESC medium comprising high-glucose DMEM (Gibco 11995065) supplemented with 15% (v/v) fetal bovine serum (HyClone SH30396.03), 1× MEM non-essential amino acids (Corning 25-025-CI), 1x Supplément GlutaMAX™ (Gibco 35050-061), 1× EmbryoMax® nucleosides (Millipore ES-008-D), 1× penicillin-streptomycin (Millipore TMS-AB2-C), 1× 2-mercaptoethanol (Gibco 21985-023), and 10³ U/mL leukemia inhibitory factor (LIF; Millipore ESG1107). The cell line was maintained at with incubation at at 37 °C and 5% CO_2_.

### 5. Immunofluorescence

All experimental steps were completed in droplets. First, mouse oocytes and embryos were fixed at room temperature in 4% PFA solution for almost 20 minutes. After washing in PBS, embryos were treated with 0.1% Triton X-100 in PBS for 20 min, blocked with 1% BSA for 1 h at room temperature, and then incubated with primary antibody (H3K4me1 and H3K27ac, 1:100 dilution) in PBS containing 1% BSA at room temperature for 1 hours or at 4°C overnight. Next the oocytes and embryos were washed and stained with Alexa Fluor 568–donkey anti-rabbit at room temperature for 1 h. Last, the samples were washed and sealed with Mounting Medium, antifading (with DAPI). In the above process, each wash needs to be repeated twice for 5 minutes each time. Finally, the original fluorescence image was obtained by Zeiss 700 confocal inverted microscope and processed by Zen (Zeiss) software. The signal intensity of the fluorescence channel 568 is adjusted to a unified parameter to ensure that the fluorescence signals between different samples can be compared.

### 6. CUT & RUN library preparation

CUT & RUN was conducted following the published protocol^[34, 35]^ with several modifications. ESCs, oocytes or embryos removed zona pellucida were transferred into 0.2 mL conventional PCR tube. The samples were resuspended by 60 μL washing buffer. Then prepare Concanavalin-coated magnetic beads 10μL for each sample were gently washed, resuspended by binding buffer and carefully added to the samples. The samples were then incubated at 23°C for 10 min on Thermomixer at 400 rpm. The samples were then held at the magnetic stand to carefully aspirate buffer, and were resuspended by 50 μL antibody buffer with the antibody H3K4me1 or H3K27ac diluted at ratio of 1:100. The samples were then incubated at 4°C on Thermomixer for overnight at 400 rpm. On the second day, the samples were held at the magnetic stand and washed by 200 μL dig washing buffer for once and 100 μL for a second time. The samples were then resuspended by 50 μL dig washing buffer with pA-MNase, and incubated at 4°C on Thermomixer for 3 h at 400 rpm. The samples were washed by 200 μL dig washing buffer for twice and 100 μL for a second time at the magnetic stand. The samples were resuspended by 100 μL dig washing buffer and balanced on ice for 2 min. Targeted digestion was performed on ice by add 2 μL 100 mM CaCl_2_ for 30 min, and reaction was stopped by adding 100 μL 2 × stop buffer and fully vortexed. The samples were then incubated at 37°C of 20 min for fragment releasing. The total samples or supernatants were digested by adding 1μL 20% SDS and 2μL Proteinase K, with 10ng carrier RNA, and the reaction was performed at 56°C for 45 min and 72°C for 20 min in PCR machine. DNA was purified by phenol chloroform followed by ethanol purification. Then purified DNA was subjected to Tru-seq library construction using NEBNext Ultra II DNA Library Prep Kit for Illumina.

### 7. CUT & RUN and ATAC-seq data processing

The paired-end CUT&RUN reads were aligned with the parameters: -t –q –N 1 –L 25 –X 1000 --no-mixed --no-discordant by Bowtie^[67]^. All unmapped reads, non-uniquely mapped reads, reads with low mapping quality (MAPQ < 20) and PCR duplicates were removed. For downstream analysis, we normalized the read counts by computing the numbers of reads per kilobase of bin per million of reads sequenced (RPKM) for 100-bp bins of the genome. To minimize the batch and cell type variation, RPKM values across whole genome were further Z-score normalized. To visualize the CUT&RUN signals in the IGV genome browser, we generated the RPKM values on a 100 bp-window base. The ATAC-seq data were mapped using mm9 (mouse) reference genome by Bowtie, the subsequent analysis is consistent with CUT&RUN data processing.

### 8. RNA-seq data processing

All RNA-seq data were mapped to mm9 reference genome by Hisat2^[68]^. The gene expression levels were calculated by Cufflinks^[69]^ (Version 2.2.1) using the refFlat database from the UCSC genome browser. For the published single cell RNA-seq data, the FPKM values of Polycomb genes from different single cells of the same developmental stages were averaged for each gene.

### 9. Peak identification and peak comparison

All CUT&RUN, ATAC-seq and RNA-seq peaks are called using MACS (Version 1.4.2)59 with nolambda -nomodel. Peaks at least 3 kb from annotated promoters (RefSeq, Ensemble, and UCSC known gene databases) were selected as distal peaks. BEDTools (version 2.27.1) was used to compare peaks.

### 10. Active, primed, poised enhancer and super-enhancer prediction

Different types of enhancers were identified based on how the three histones are combined. The combination of H3K4me1 and H3K27ac was identified as active enhancer, whereas the combination H3K4me1 and H3K27me3 was regarded as poised enhancer. Besides, the H3K4me1 only was the primed enhancer^[21, 22]^.

For super-enhancer, we used sequencing data from H3K27ac to identify super-enhancers at various stages of preimplantation embryos. MACS 1.4.2 determines the enrichment area of H3K27ac in each stage by parameters’ -p 1E-9 ‘, ‘-keep-dup = auto’, ‘-W-s-space = 50’, ‘-g hs’. Using the output from MACS, ROSE identified the super-enhancer. The constituent enhancers were stitched together within 12500 bp. If the constituent enhancer is completely contained within the promoter region (window ± 2000 bp from the transcription start site), it is excluded from splicing. Finally, we separate the super-enhancer from the typical enhancer by separating the H3K27ac signal from an inflection point in the enhancer class.

### 11. Motif analysis

Enrichment of known motifs within promoter and enhancer regions was analyzed with HOMER with default parameters and a fragment size of 200 bp. All known motifs used in our study were defined by HOMER^[70]^.

## Credit author statement

Kaiyue Hu: Data curation, Writing-original draft, Visualization, Software. Chunling Wang: Writing - review & editing, Validation. Dong Fang: Methodology, Visualization. Jiacheng Lu: Methodology, Visualization. Xiangrui Meng: Methodology, Visualization. Lingling Chen: Methodology. Yage Yao: Methodology. Jia Guo: Methodology. Sarmir Khan: Methodology. Wenbo Li: Methodology. Yaqian Wang: Methodology. Yang li: Methodology. Hao Chen: Supervision, Resources. Jiawei Xu: Supervision, Resources.

## Funding

This study was supported by the National Natural Science Foundation of China (31870817 and 32170819), the National Key R&D Program of China (2019YFA0110900 and 2019YFA0802200), the Science and Natural Science Foundation of Henan Province (22100026/20), the Special Funding of the Henan Health Science and Technology Innovation Talent Project, the Henan Provincial Medical Science and Technology Research Program (SBGJ202402099) and the Natural Science Foundation of Henan Province (252300423781).

## Acknowledgments

We thank all participants for the work, members of Jiawei Xu and Hao Chen for the research design. Other authors contributed to manuscript revision, read, and all the authors approved the submitted version. Specially, we thank a lot for the help revising and advising of Bofeng Liu, TsingHua University.

## Data Availability Statement

The data that support the findings of this study are available from the corresponding author upon reasonable request.

## Conflict of Interest

The authors declare no conflict of interest.

## Notes

### Competing Interest Statement

The authors have declared no competing interest.

### Summary of Updates

This version has been revised to improve the clarity and readability of the text.

