## supplementary Figures for "Histone modifications analysis reveals enhancers reprogramming during maternal-to-zygotic transition"

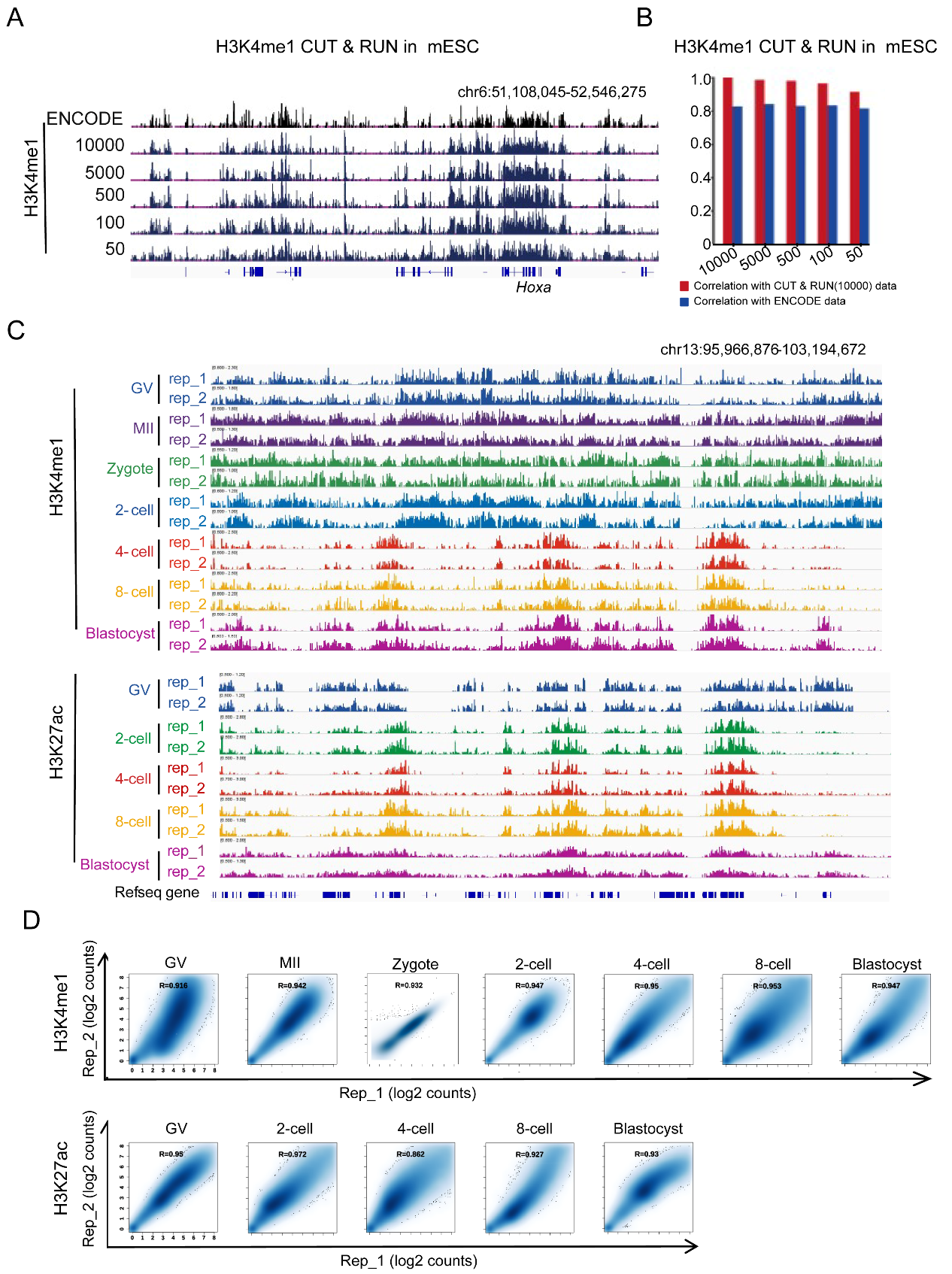

Fig. S1. Validation of H3K27ac CUT & RUN data in mouse oocytes and early embryos. A, IGV browser showing the H3K4me1 signals detected in mouse embryonic stem cells with different cell concentration gradients and the H3K4me1 generated from encode data. B, Correlation analysis of H3K4me1 CUT & RUN data in mouse embryonic stem cells. C, IGV browser showing two replicates of H3K4me1 and H3K27ac enrichment in oocytes and preimplantation embryos. D, Correlation analysis of the two replicates of H3K4me1 and H3K27ac CUT & RUN data in various stages.
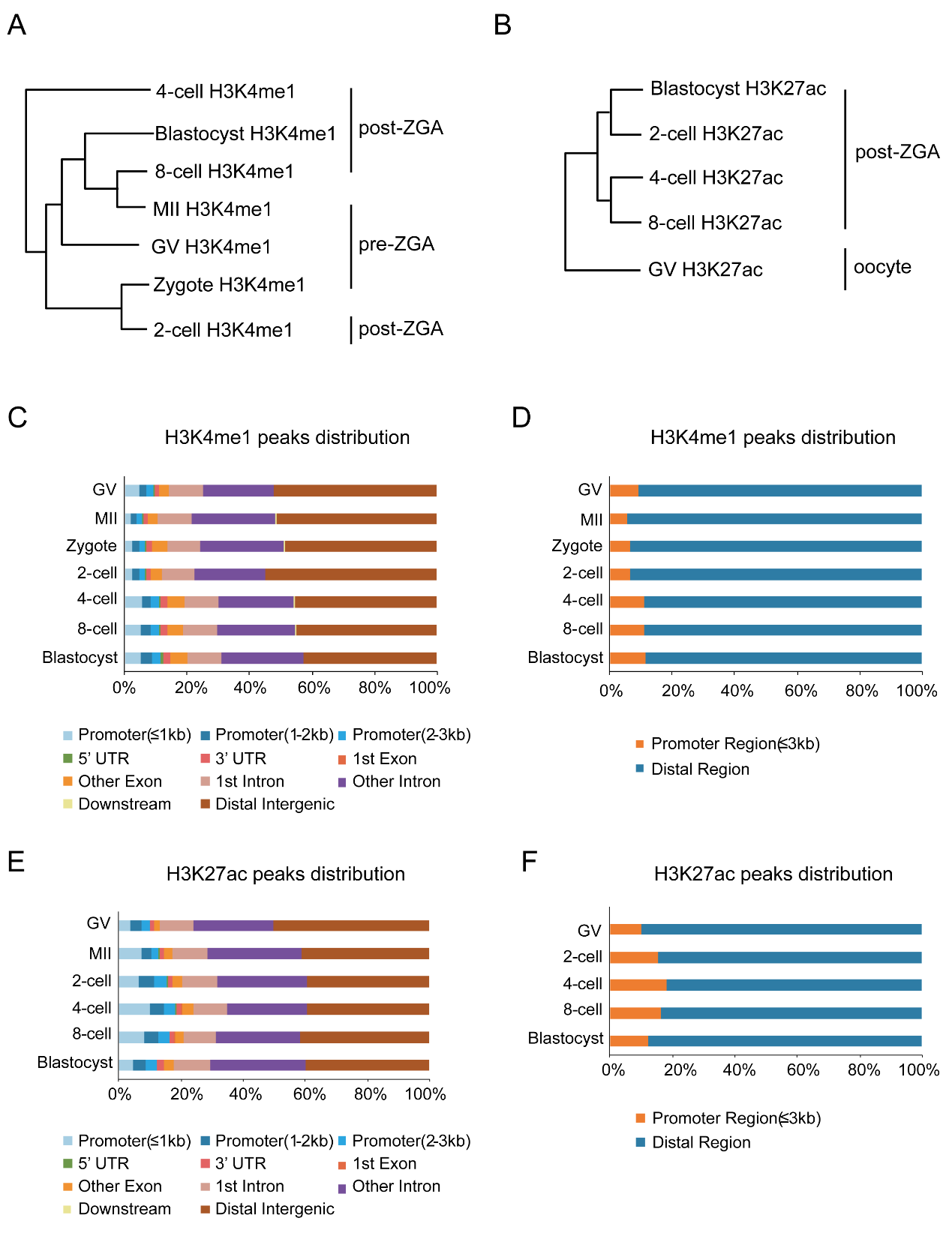

Fig. S2. The distribution characteristics of H3K4me1 and H3K27ac across wide genome. A, Hierarchical clustering of global H3K4me1 enrichment. B, Hierarchical clustering of global H3K27ac enrichment. C, The distribution of H3K4me1 in different genome elements. D, The distribution of H3K4me1 in promoter region and distal region. E, The distribution of H3K27ac in different genome elements. F, The distribution of H3K27ac in promoter region and distal region.

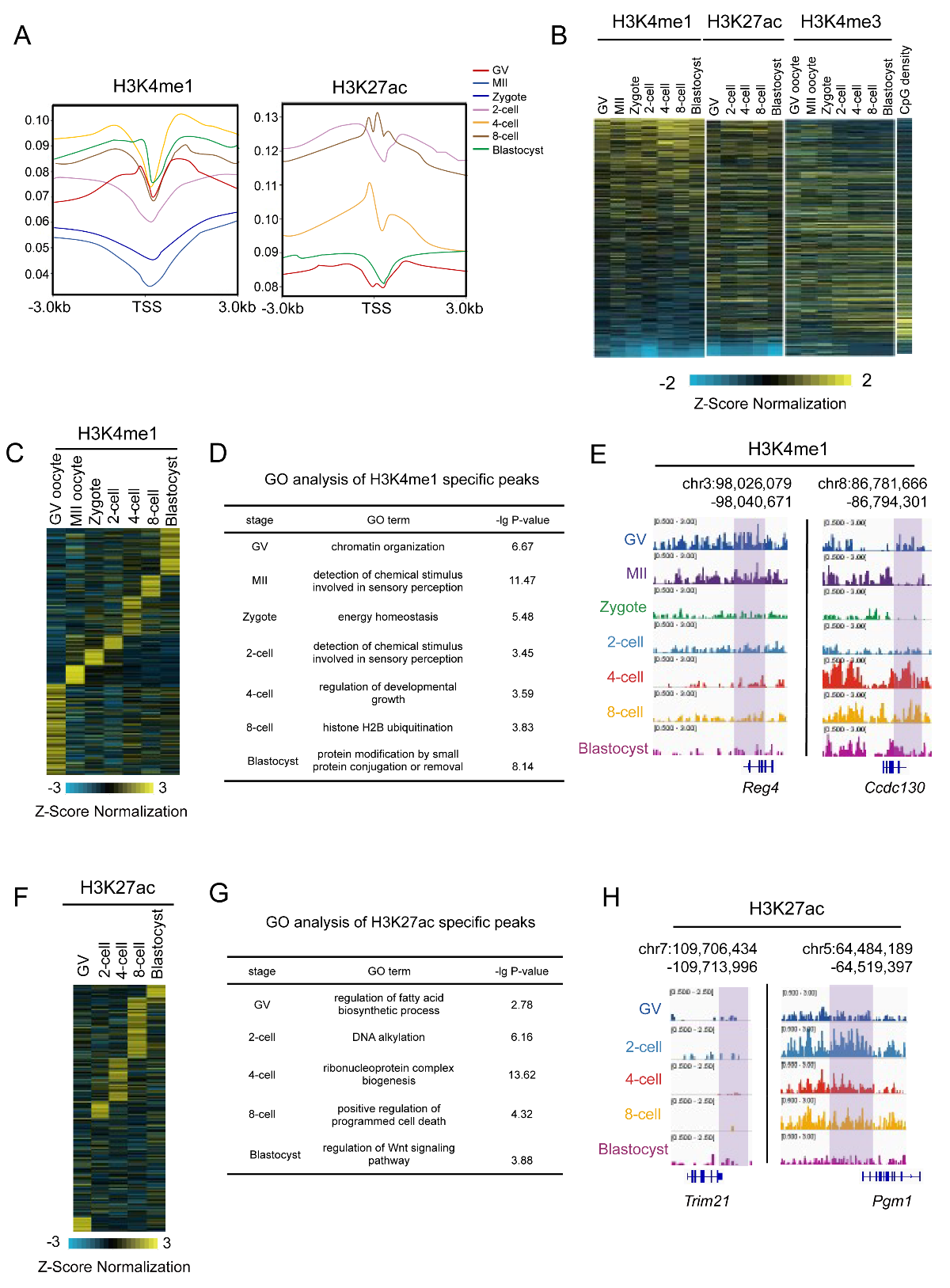

Fig. S3. H3K4me1 and H3K27ac distribution in promoter regions. A, The distribution pattern of H3K4me1 andH3K27ac in promoter regions. B, Heatmap showing H3K4me3,H3K4me1,H3K27ac and CpG density signals in promoter regions. C, The stage-specific peaks of H3K4me1 in promoter regions. D, The GO analysis related to the stage-specific peaks identified in Fig. S3C. E, The representative loci with the stage-specific H3K4me1 distributed. F, The stage-specific peaks of H3K27ac in promoter regions. G, The GO analysis related to the stage-specific peaks identified in Fig. S3F. H, The representative loci of the stage-specific H3K27ac distribution.

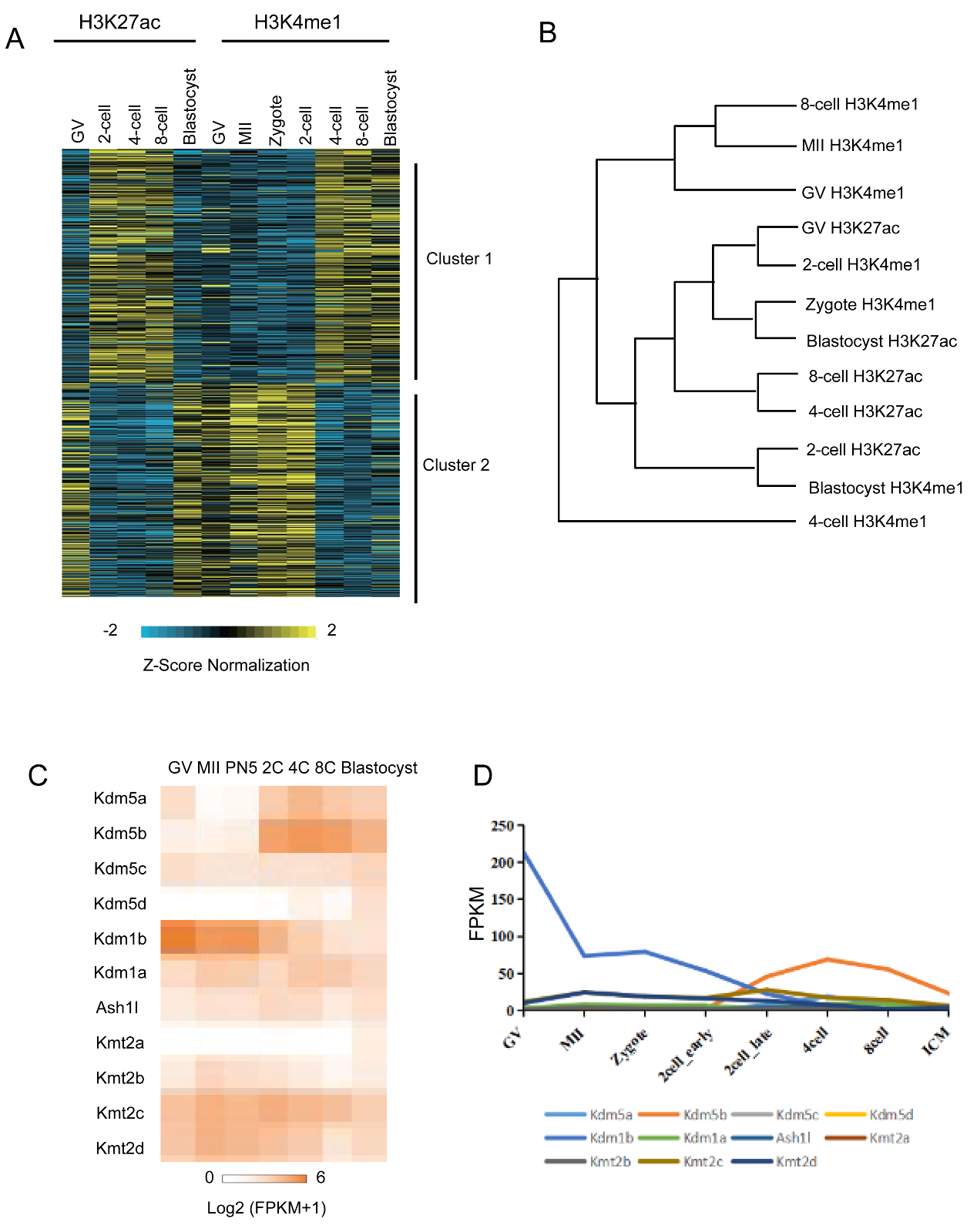

Fig. S4. The reprogramming of H3K4me1 and H3K27ac in distal regions and the related enzymes. A, K-means cluster analysis of H3K4me1, and H3K27ac enrichment in distal regions. B, Hierarchical clustering of H3K4me1 and H3K27ac enrichment in distal regions. C, Heatmap showing the expression level of the mRNA for H3K4-related enzyme. D, Line chart showing the expression level of the mRNA for H3K4-related enzyme.

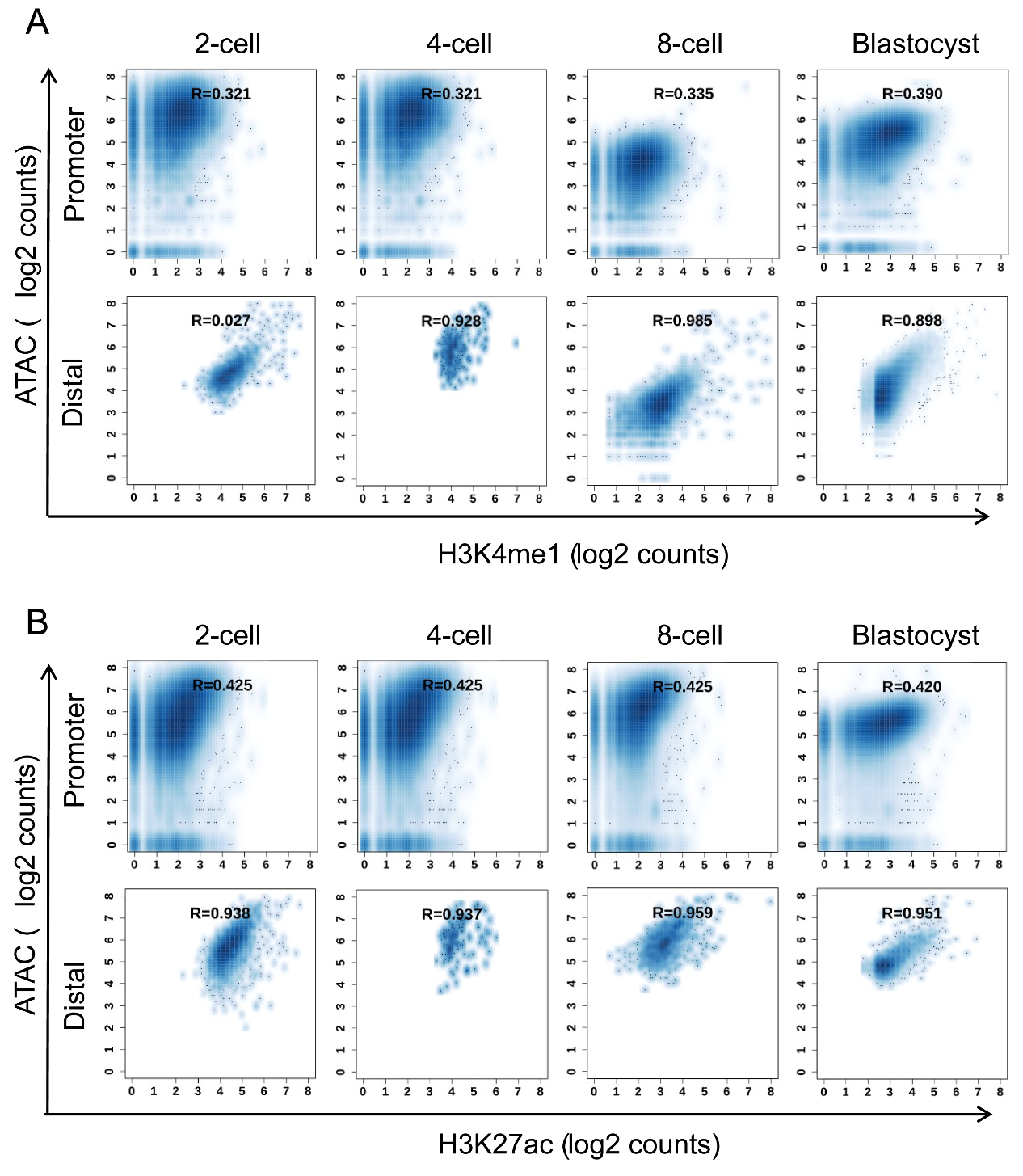

Fig. S5. Correlation analysis of H3K4me1 and H3K27ac to ATAC-seq data. A, Correlation analysis of H3K4me1 and ATAC-seq in promoter region and distal region. B, Correlation analysis of H3K27ac and ATAC-seq in promoter region and distal region.

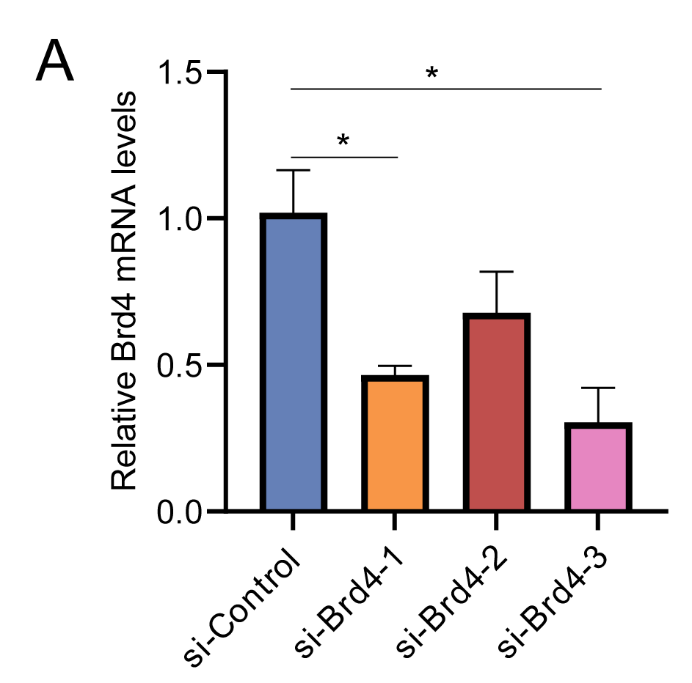

Fig. S6. Screening of siRNA targets for Brd4 knockdown in embryonic stem cells. A, The relative Brd4 mRNA levels in embryonic stem cells transfected with si-Control and three different siRNAs targeting Brd. Data are presented as mean ± SEM. **P* < 0.05.
